# Comparative Analysis of Linear and Nonlinear Dimension Reduction Techniques on Mass Cytometry Data

**DOI:** 10.1101/273862

**Authors:** Anna Konstorum, Nathan Jekel, Emily Vidal, Reinhard Laubenbacher

## Abstract

Mass cytometry, also known as CyTOF, is a newly developed technology for quantification and classification of immune cells that can allow for analysis of over three dozen protein markers per cell. The high dimensional data that is generated requires innovative methods for analysis and visualization. We conducted a comparative analysis of four dimension reduction techniques – principal component analysis (PCA), isometric feature mapping (Isomap), t-distributed stochastic neighbor embedding (t-SNE), and Diffusion Maps by implementing them on benchmark mass cytometry data sets. We compare the results of these reductions using computation time, residual variance, a newly developed comparison metric we term neighborhood proportion error (NPE), and two-dimensional visualizations. We find that t-SNE and Diffusion Maps are the two most effective methods for preserving relationships of interest among cells and providing informative visualizations. In low dimensional embeddings, t-SNE exhibits well-defined phenotypic clustering. Additionally, Diffusion Maps can represent cell differentiation pathways with long projections along each diffusion component. We thus recommend a complementary approach using t-SNE and Diffusion Maps in order to extract diverse and informative cell relationship information in a two-dimensional setting from CyTOF data.

## 1 Introduction

Many current questions in the field of immunology require cell classification to analyze the responses of immune cells to external stimulants [21]. Flow cytometry, which allows the single-cell analysis of marker expression via fluorescent-antibody labeling, has been used by immunologists to classify and sort cells since the 1960’s [35]. Mass cytometry, also known as CyTOF (cytometry by time of flight), is a newly developed technique for single cell measurement which has opened the door to new functional and phenotypic information that can help immunologists in these analyses. Unlike flow cytometry, where the number of markers analyzed is lim-ited by fluorescence spectral overlap, in mass cytometry antibodies are labeled with heavy metal ion tags that can be read using inductively coupled plasma mass spectrometry (ICP-MS) [15]. Hence, a main advantage of CyTOF over flow cytometry is its ability to measure over three dozen biomarkers simultaneously [8]. While this capability is a breakthrough in the observation of single cell parameters, it comes with its own set of challenges. As with any high throughput technique, mass cytometry produces such large amounts of data that it is difficult to extract the salient information. For the data to be useful, it is necessary to apply innovative data analysis techniques [4].

One of the most powerful tools for analyzing high dimensional data is dimension reduction. Dimension reduction renders high dimensional data in a few dimensions while preserving its significant characteristics. There exists a wide array of techniques for dimension reduction, each of which aims to preserve a specific feature of the data by finding the intrinsic degrees of freedom with respect to that feature. While marker-specific information is lost in this process, critical information regarding intercellular relationships is extant in dimension-reduced data.

In order to decide which dimension reduction technique to use, it can be helpful to know how each technique performs on mass cytometry data with respect to a given criterion. We thus consider how each of four popular dimension reduction techniques – principal component analysis (PCA), t-distributed stochastic neighbor embedding (t-SNE), isometric feature mapping (Isomap), and Diffusion Maps, perform on manually gated benchmark mass cytometry datasets with respect to computation time, neighborhood proportion error, residual variance, and ability to cluster known cell types and track differentiation trajectories. We choose these four dimension reduction methods for the following reasons: PCA is a highly-utilized and computationally efficient linear method for dimension reduction, t-SNE exaggerates cluster formation and has been extensively employed in CyTOF analysis via the viSNE toolkit [2]; Isomap, on the other hand, does not exaggerate cluster formation, and preserves geodesic distances between points that can allow for preservation of nonlinear interactions (unlike PCA) and global relationships between cells and cell clusters (unlike t-SNE). Finally, Diffusion Maps, like Isomap, can preserve nonlinear interactions and global relationships between cells, and has the additional property of being highly suitable to extract differentiation trajectories between cells [20], which is often of interest in experimental studies utilizing CyTOF.

In addition to dimension-reduction techniques, there exist several computational algorithms for analysis of mass cytometry data, including but not limited to methods based on agglomerative clustering (SPADE [33], FLOW-MAP [47], and Citrus), neural networks (FLOW-SOM [43]), and graph-based, trajectory-seeking, algorithms (Wanderlust [7], Wishbone [37]) (see [10, 29, 36] for recent reviews of the applications of these and other algorithms to mass cytometry). The SPADE and FLOW-SOM algorithms organize clusters of points into a minimal spanning tree (MST), while the FLOW-MAP uses a highly connected graph structure to connect clusters instead. The assumption that all clusters are connected can create artificial relationships between clusters and/or cells if the common progenitor of two cell types is not included in the population. Similarly, the assumption by trajectory-seeking algorithms such as Wanderlust (which does not allow branching) or Wishbone (which does), that all cells are part of a developmental hierarchy may not hold for more heterogeneous cell populations. We focus on dimensionality-reduction methods since they require fewer assumptions about the nature of cell-cell relationships for any given analysis, and hence provide the most unsupervised analysis possible for this type of data.

## 2 Methods

### 2.1 Dimension Reduction Techniques

We provide a methodological overview of the dimension reductions techniques that we utilize in our comparative analysis. As the preponderance of such techniques grows, it is imperative that researchers and end users maintain a firm grasp on their underlying technical frameworks. With the summaries of the techniques below, we hope to aid in this goal.

#### 2.1.1 Principal Component Analysis (PCA)

PCA is a linear dimension reduction method that is designed to preserve the variability of a data set, and has been recognized as an effective method to analyze high-throughput biological data [34]. PCA has been used on CyTOF data in a variety of settings, including a demonstration that different subsets of CD8+ T cells actually form a continuum, with cells that can be considered intermediate phenotypes bridging different subsets [30], to help develop a T cell epitope binding prediction algorithm [31], and to show that a conserved T cell transcriptional profile including CD161 expression exists among different T cell subtypes [17].

Given a data matrix *X_m×n_*, with *m* attributes and n samples, it’s covariance matrix *C_x_* is given by

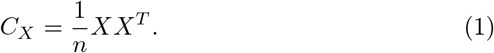

The diagonal elements of the matrix *C_X_* represent the variance of the attributes, and the off-diagonal elements the covariance. Principal component analysis seeks a transformation

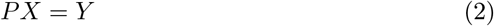

such that the diagonal elements of *C_Y_* are rank-ordered and the off-diagonal elements equal to zero. It can be shown that this *P* is the eigenvector matrix of *C_X_*\*, where *X*\* is the mean-centered matrix of *X*. Each of these eigenvectors, termed principal components, are linear combinations of the original attributes that are orthogonal to each other. The first principal component comprises the most variance of the data that can be captured by a linear combination of the attributes, the second principal component comprises the most variance of the data that can be captured by a linear combination of the attributes after the first principal component has been accounted for, and so on. In practice, the principal components are often calculated using the singular value decomposition (SVD), which can take any matrix *X_m×n_* and convert it into a product of an *m × m* orthogonal matrix (the left singular vectors of *X*), an *m × n* rectangular diagonal matrix (containing the singular values of *X*), and an *n × n* orthogonal matrix (containing the right singular vectors of *X*). The principal components of *X* are the right singular vectors of *X*\*, and since calculating the SVD of a matrix *X* is more computationally efficient than finding the eigenvectors of its covariance matrix, it is commonly used to determine the principal components of *X* in practice. For more details regarding the technical background of PCA and SVD, see the tutorial by Shlens [39].

#### 2.1.2 T-distributed Stochastic Neighbor Embedding (t-SNE)

The t-SNE algorithm is a non-linear dimension reduction method that associates probability distributions to all high-dimensional points and seeks to maintain a probability distribution profile in the low-dimensional embedding. It has been used to visualize and cluster data in a variety of CyTOF-related studies, including healthy and leukemic bone marrow [2], the human mucosal immune system [44], and phenotypic diversity of human regulatory T cells (Tregs) [27]. T-SNE is the reduction technique behind the viSNE algorithm for mass cytometry data [2].

In t-SNE, the similarity *p_ij_* between two points *i* and *j* is calculated using a Gaussian joint probability distribution, which gives a joint probability distribution *P* = {*p_ij_* }. To compute the joint probability distribution, *Q* = {*q_ij_*}, in the low-dimensional space, a Cauchy distribution is used since it has a heavier tail than the Gaussian distribution and can thus mitigate the effect of the ‘crowding problem’, which can push points at moderate distances too far away in the low-dimensional map, and points that are nearby too close together [26]. The goal of t-SNE is to minimize the difference between the two distributions, hence a measure for the divergence of two probability distributions, the Kullback-Leibler (KL) divergence, is minimized using gradient descent in the cost function

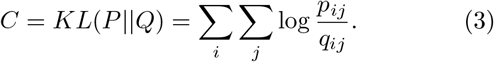

T-SNE is effective at extracting natural clusters since data points further apart from each other in the high-dimensional space are assigned disproportionately smaller probabilities, which exaggerates the boundaries between clusters in the low dimensional embedding [26].

#### 2.1.3 Isometric Feature Mapping (Isomap)

Isomap is another well-known nonlinear dimension reduction method [42]. Unlike t-SNE, it does not exaggerate distances between clusters, hence can be used to obtain more appropriate distance measures between different cell types and to investigate differentiation trajectories, which do not naturally lend themselves to clustering. Becher et al. [5] used Isomap to investigate the relationships between different myeloid subsets based on CyTOF data, and identified potential differentiation trajectories. Notably, they used it as an adjunct to t-SNE, which was used to aid in clustering different cell types in two dimensions, but was identified by the authors as not an appropriate method to identify differentiating cells. Similarly, Wong et al. [46] used Isomap to probe differentiation of CD4^+^ T cells in peripheral tissue (blood and tonsils) and in conjunction with t-SNE, identified clusters of cells representing increasingly differentiated states along the trajectories. Using three different microarray datasets, Dawson et al. [13] were able to show that Isomap was able to effectively cluster and establish relationships between different treatments for spinal cord injury data, gene expression data from different rat tissues, and a high-throughput drug screen against acute myeloid leukemia (AML), showing the effectiveness of Isomap to distinguish important phenotypes amongst a diverse set of datasets.

In Isomap, the distance between two points *i* and *j*, *δ_i,j_* in the original data is measured by the *geodesic distance*: for a point *x* in the original dataset *X*, points *j* within *ϵ* Euclidean distance of *i*, or the *K*-nearest neighbors of *i*, are assigned their Euclidean distance to the entry *δ_i,j_* in the geodesic distance matrix *D_G_* = {*d*_*ij*_}. For points *j* outside of a neighborhood of *i, δ_i,j_* is assigned the shortest path distance between *i* and *j*. Isomap seeks to preserve the geodesic distances recorded in *D_G_* for all points in *X* by using the linear dimension-reduction technique multidimensional scaling (MDS) on *D_G_* [12, 18]. MDS can identify a low-dimensional representation of *X, Y*, such that the error function

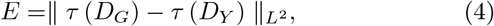

is minimized, where || · ||_*L*^2^_ is the *L*^2^ norm, *D_Y_* the matrix of Euclidean distances of points in *Y*, and *τ* is the transformation

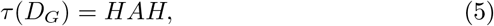

where *H* = *I_n_* – (1/*n*) is a ‘centering matrix’, *n* the number of data points, and 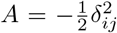. The *τ* operator allows recovery of *Y* via the eigendecomposition of *τ*(*D_G_*).

#### 2.1.4 Diffusion Maps

Diffusion Maps is based on a spectral analysis of a diffusion matrix based on random walk probabilities between cells (see below), and has been championed for use in differentiation analysis of single-cell data, especially as it offers a robustness to noise that Isomap lacks [11]. For example, Haghverdi et al. [20] used Diffusion Maps to show that for both a toy differentiation dataset and qPCR and RNA-Seq data from differentiation processes including mouse hematopoietic differentiation, differentiating cells from mouse zygote to blastocyst, and human preimplantation embryos, that Diffusion Maps can identify differentiation trajectories on a low-dimensional representation that is robust to noise and sampling heterogeneity.

We base our discussion of Diffusion Maps on [14]. If we take *p*(*x,y*) as the probability of jumping, in a random walk, between two points *x* and *y* in a dataset X, then we can calculate *p* via a symmetric and positivity-preserving Guassian kernel, *k*(*x,y*). We take the diffusion matrix *P* = {*p*(*x_i_,x_j_*)}, with *x_i_,x_j_* ∈ *X*. Note that powers of *P, P*^*t*^ = {*p_t_*(*x_i_,x_j_*)} give probabilities of moving from *x_i_* to *x_j_* in *t* steps. The diffusion distance is then taken to be

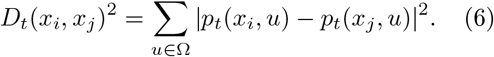

A large number of high probability paths between two points *x_i_* and *x_j_* will give a small *D_t_*(*x_i_,x_j_*), and vice-versa. Diffusion Maps seeks to identify a low-dimensional embedding *Y* of the points in X such the Euclidean distance between two points *y_i_,y_j_* ∈ *Y* mapped from *x_i_* and *x_j_* will approximate the diffusion distance *D_t_*(*x_i_,x_j_*). The lowdimensional embedding *Y* is found by assigning to Y the dominant eigenvectors associated with a spectral decomposition of P [11, 14].

### 2.2 Benchmark Data

We consider two manually-gated benchmark datasets to compare the dimension reduction techniques. The availability of manual gating data alongside markers for the cells allows for a comparison of how the dimension reduction techniques can identify known cell populations and the relationships between the known populations. The first dataset was generated by Bendall et al. [9] and consists of ~170,000 cells with 13 markers, of which ~50% have been manually gated into 24 cell populations. The cells are derived from healthy primary human bone marrow (bone marrow mononuclear cells, BMMCs), and hence represent the hematopoietic differentiation spectrum in the bone marrow (Figure 1). We repeat our analysis on an additional benchmark BMMC dataset from two healthy human donors generated by Levine et al. [24] that consists of ~160,000 cells with 32 markers, of which ~35% have been manually gated into 14 cell populations. Both datasets were obtained from [1] Additional information on the benchmark datasets and the analysis of the 32 marker dataset is found in the Supplementary Material.

**Figure 1:**
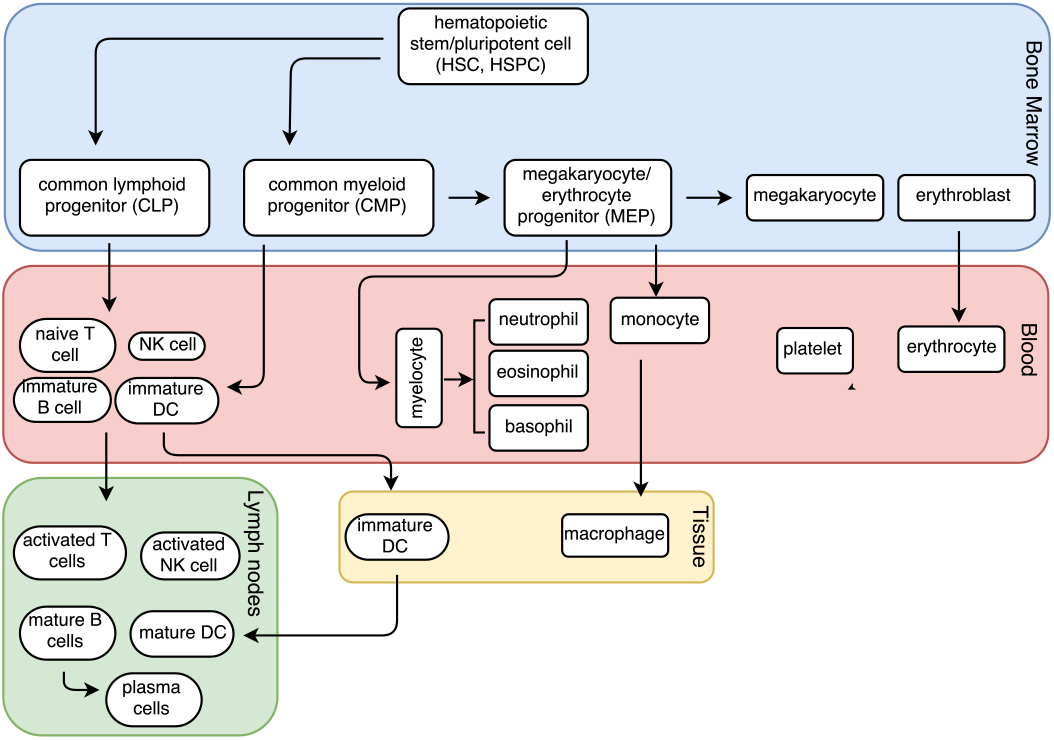
Differentiation of cells from hematopoietic stem cells. Cells from different stages in the hematopoietic differentiation process were collected in both the 13- and 32-marker benchmark datasets. Figure adapted from [28].

### 2.3 Comparison Metrics

#### 2.3.1 Computation Time

The computation time for each method depends heavily on the software and the implementation used. For each technique, we chose the most optimized implementation currently available for R [41], (Table 1). All dimension reductions were executed on a MacBook Air (2.2GHz Intel Core i7 processor and 8 GB memory). Currently, the most widely used dimension reduction technique for mass cytometry data is t-SNE, which is the method behind the popular viSNE algorithm. ViSNE takes a random sample of between 6,000 and 12,000 cells to perform its reductions [2]. We chose to take three random subsets of the data, each consisting of 10,000 cells, and measured the run-time for one reduction from *n* = 13 to *m* = 2 dimensions on each subset. Our final computation time result is the average of the three reductions for each technique.

**Table 1:** The R packages used to implement each dimension reduction technique, as well as the repositories from which they are available [32, 3, 22, 23].

| Method | R Package | Repository |
| --- | --- | --- |
| PCA | stats (princomp) | R |
| t-SNE | Rtsne | CRAN |
| Isomap | vegan | CRAN |
| Diffusion Maps | destiny | Bioconductor |

T-SNE is not designed to operate efficiently on more than 10,000 cells, although a random walk based version has been proposed that will reduce a subset of data while using the information from the entire data set [26]. The *Rtsne* package is a version of the Barnes-Hut implementation of t-SNE, which is an accelerated implementation and the best suited for large data sets [25]. The *vegan* package provides a traditional implementation of the Isomap algorithm, which involves complex computations that render it highly inefficient on large data sets. The most notable inefficiencies in the algorithm are the construction of neighborhood graphs and the MDS eigenvalue decomposition. A new variation of Isomap called L-Isomap uses landmark points to calculate distances providing computational simplification [40]. Due to potential topological instability exacerbated by the landmark approach, questions of how to choose the landmark points [38], and a lack of an implementation in R, we do not use L-Isomap in our analysis, although it can be considered for future applications to CyTOF. The *destiny* package for R provides an implementation of Diffusion Maps developed by Angerer et al. [3] specifically for high-throughput single-cell data. Parameters used for each dimension reduction method employed in R are summarized in Supplementary Table S3.

#### 2.3.2 Neighborhood Proportion Error (NPE)

An important motivation for mass cytometry data is the clustering of cells by phenotype. The cells in the benchmark data set have been manually assigned to subtypes based on their marker expression. The degree to which the cells cluster together in the input space is a highly informative characteristic of any mass cytometry data set. Using the subtype assignments provided by manual gating, we developed the NPE to measure how effectively each dimension reduction technique translates the cell proximities within a subtype from the input space to the low-dimensional embeddings (Figure 2). Unlike the neighborhood trustworthiness measure proposed by Venna and Kaski [45], which measures the degree of overlap between the original and projected neighborhood, the NPE seeks to ascertain whether the projection maintains neighborhood (dis)similarity as follows: if a subtype of cells, *s*, does not cluster closely in the original space, i.e. cells of subtype *s* have a low proportion of neighbors of subtype *s*, then the dimension reduction should show a similarly low proportion of neighbors of like subtype for cells of subtype *s*.

**Figure 2:**
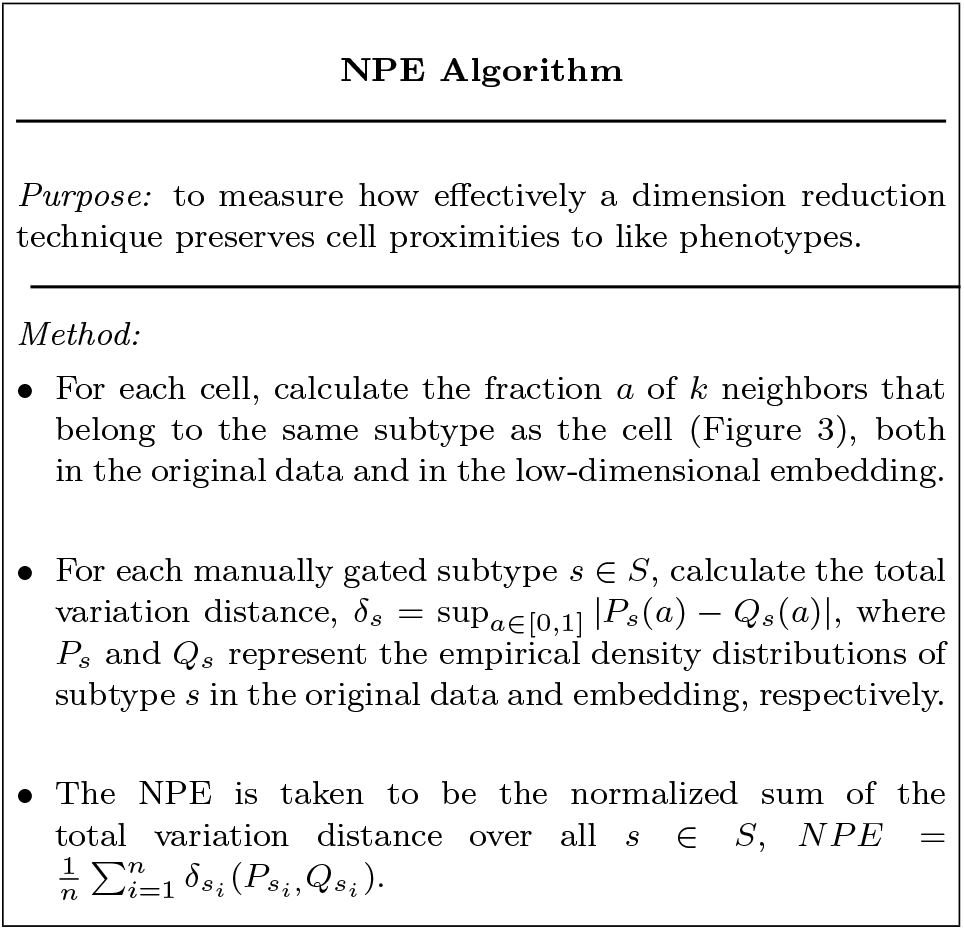
Overview of the Neighborhood Proportion Error (NPE) algorithm.

In the NPE calculation, all columns (which represent the markers measured in the experiment) are first normalized to mean 0 and variance 1. Every data point (or cell) is then assigned a neighborhood of the *k* closest points, as measured using the Euclidean distance metric. For each neighborhood, the fraction, *a*, of neighbors that belong to the subtype of the cell is then calculated (Figure 3).

**Figure 3:**
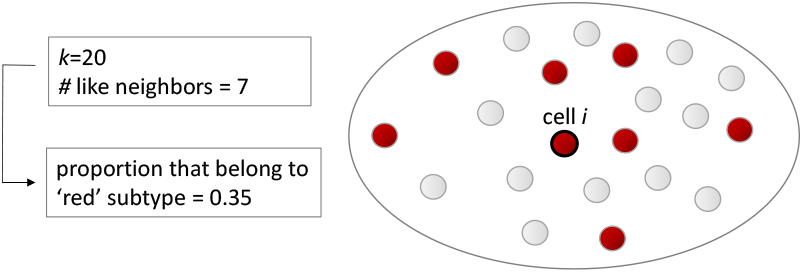
A sample neighborhood of *k* = 20 points showing the fraction of cells that are of the same sub-type as cell *i* (black outline).

The results are converted to an empirical density estimate *P_s_*(*a*) in the original space and *Q_s_*(*a*) in the dimension-reduced space, for each subtypes *s* ∈ *S*, where *S* is the set of all manually gated subtypes. The error between the density estimates for each *s* is calculated using total variation distance [19],

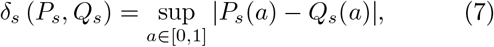

where *a* represents the fraction of like neighbors that constitutes the domain of *P_s_* and *Q_s_*. The total variation distance, also known as the “statistical distance” is a measure of the largest difference between the probabilities that *P_s_* and *Q_s_* can assign to the same event, *s*. The NPE is calculated as the normalized sum of *δ_s_* over all the subtypes *s_i_* ∈ *S*,

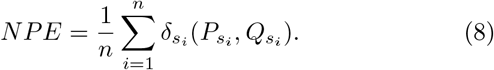

#### 2.3.3 Residual Variance

In addition to the measure of local preservation that NPE provides, we also want to compare our methods with respect to global error. Residual variance gives a measure of the global variance not accounted for by the dimension reduction. It is defined as [42]

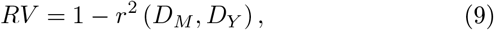

where *r* is the Pearson correlation coefficient,

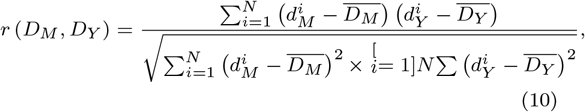

and *D_M_* and *D_Y_* are the respective high- and lowdimensional distance matrices. *D_Y_* consists of the pairwise Euclidean distances between the points in the reduced space; however, *D_M_* is different for each method. In PCA, *D_M_* is composed of the pairwise Euclidean distances in the input space. In Isomap, *D_M_* consists of the geodesic distances derived from the weighted graph *D_G_* (Section 2.1.3). In Diffusion Maps, *D_M_* = *D_t_*, where *D_t_* is defined in Equation (6). Because the cost function for t-SNE (Equation (3)) is nonlinear, the relationships between the input and output probability distributions, *P* and *Q,* respectively, are nonlinear (Section 2.1.2) and the Pearson correlation coefficient, which measures the linear correlation between two variables, will not give a proper measure of the information preserved in the dimension reduction, thus residual variance was not calculated for t-SNE. For the other methods, residual variance was calculated for dimension reductions for low-dimensional projections of dimension *m* =1, 2, …, 7.

#### 2.3.4 Visualization: clustering and differentiation

Since an important goal of all the dimension reduction methods is visualization and identification of known and new cell subtypes, as well as the tracking differentiation trajectories, we visualized the two dimensional reductions using a color overlay of the gated cell subtypes, and considered whether (1) manually gated subtypes clustered together in the visualizations and (2) the dimension reduction methods were able to capture known differentiation trajectories.

## 3 Results

The results presented are from application of the comparison metrics to one of the 10K subsets of the 13-dimensional dataset. Results for the other 10K subsets of the 13-dimensional dataset, as well as for a 10K subset of the 32-dimensional dataset, are found in the Supplementary Material, Sections S3-S4. We summarize the results in Table 2.

**Table 2:** Summary of performance for the dimension reduction methods on the 13 marker dataset based on the various comparison metrics. Computation time, NPE, and RV results show averages over the three data subsets. NPE: neighborhood proportion error, RV: residual variance.

| Comparison Metric | Dimension Reduction Method |  |  |  |
| --- | --- | --- | --- | --- |
|  | *PCA* | *Isomap* | *t-SNE* | *Diffusion Maps* |
| *Comp. Time (m=2)* | 0.034s | 11428.42s | 104.174s | 122.676s |
| *NPE (m=2, k=20)* | 0.68 | 0.64 | 0.38 | 0.44 |
| *RV (m=2)* | 0.230 | 0.451 | N/A | 0.916 |
| *Visualization* | Cell subtypes difficult to distinguish, no discernible differentiation trajectories. | Cell subtypes difficult to distinguish, no discernible differentiation trajectories. | Different cell types easy to distinguish, no discernible differentiation trajectories. | Different cell types can sometimes be distinguished, discernible differentiation trajectories. |

### 3.1 Computation Time

Computation time calculations show pronounced differences between the reduction methods (Figure 4). Isomap is the slowest by a wide margin, requiring approximately 3 hours. The other three methods proved more practical, with t-SNE and Diffusion Maps requiring 1.5-2 minutes, and PCA less than one second.

**Figure 4:**
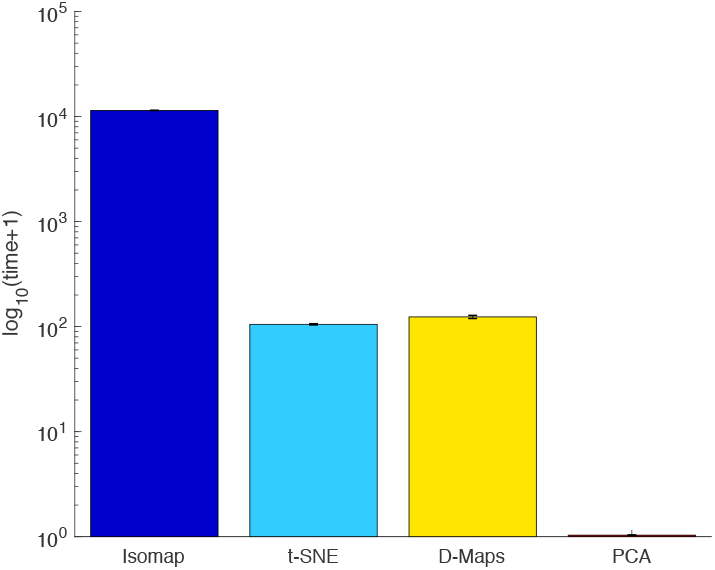
Dimension reduction time for three random subsets of 10,000 cells from *m* =13 dimensions to *n* = 2 from the benchmark dataset. Results are plotted using a semilogarithmic plot in increasing order of efficiency.

### 3.2 Neighborhood Proportion Error

NPE was calculated for two-dimensional reductions of all tested methods (Figure 5). We observe that as the neighborhood size increases, NPE values begin to stabilize and, most importantly, do not change rank order with respect to the dimension reduction methods, thus showing a robustness in results to k. PCA shows the most error, indicating that its neighborhood proportions with respect to cell subtype were the most altered through the process of dimension reduction. T-SNE and Diffusion Maps preserve neighborhood proportion significantly better than Isomap and PCA, signaling that these methods are better able to preserve the local structure and neighborhood proportions of gated populations, which may be important when considering inherent relationships between cells of known phenotypes vis-a-vis their CyTOF profiles.

**Figure 5:**
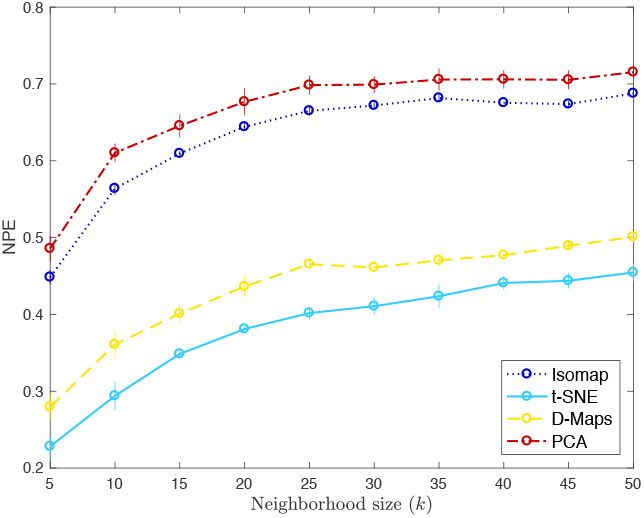
Neighborhood Proportion Error (NPE) as a function of *k*, the size of the neighborhood, and the dimension reduction method (PCA, t-SNE, Isomap, and Diffusion Maps).

### 3.3 Residual Variance

Residual variance (RV) was calculated for PCA, Isomap, and Diffusion Maps for dimension reductions to *m* = 1, 2, …, 7 dimensions (Figure 6). We observe that PCA and Isomap show a strong reduction in RV up to *m* = 3, after which the reductions are more modest. This ‘elbow effect’ can signal that the intrinsic dimensionality (*m*) of the data is between 3 and 4 [42]. Surprisingly, Diffusion Maps, although a strong performer in the NPE method, shows a higher RV at every *m* than the other methods, and does not display an elbow effect. This is not necessarily an outcome of properties of the methods themselves, since Diffusion Maps was found to have a lower RV than Isomap and PCA at higher dimensions in a comparative analysis of dimension reduction techniques for the free energy landscapes of peptide folding [16]. Although these results show that the intrinsic dimensionality of the dataset may be higher than *m* = 2, we proceed with our comparative analysis to reductions of two dimensions since this is the dimensionality at which the majority of users are interested in considering the data.

**Figure 6:**
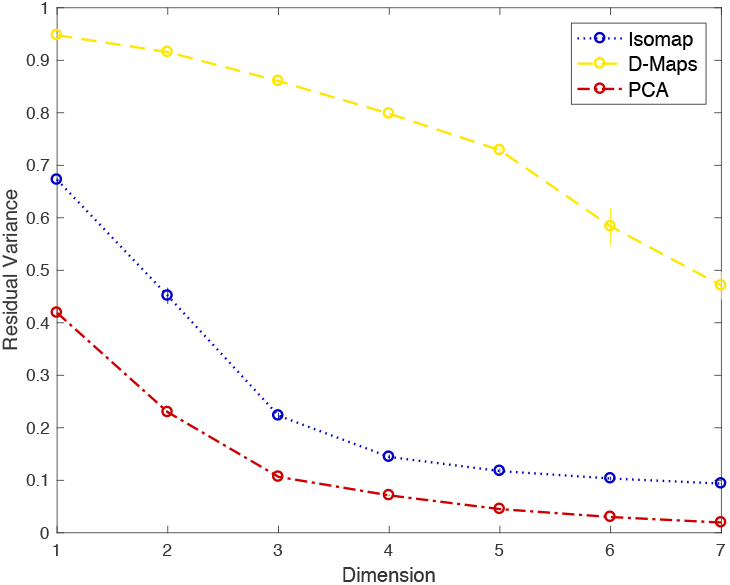
Residual variance error for Isomap, Diffusion Maps, and PCA for low-dimensional embeddings of dimension *m* =1, …, 7.

### 3.4 Visualization: clustering and differentiation

There are two specific phenomena that experimentalists are generally interested in observing in the two-dimensional embeddings: phenotypic clusters and differentiation trajectories. Phenotypic clustering refers to cells with the same manually-gated subtypes occupying a region of the plot near to each other, preferably to a degree that allows distinct populations with clear boundaries to be identified. Differentiation trajectories refer to cell subtypes that are arranged along a clear path in the order that cell differentiation occurs.

The visualizations show that the different dimension reduction techniques differ in their ability to identify phenotypic clusters and differentiation trajectories (Figure 7). PCA does not demonstrate clear separation of cell subtypes, there is a significant mixing between the different populations, making this embedding ineffective for subtype classification. PCA also does not produce any clear paths along which we can observe the process of cell differentiation. Isomap shows similarly extensive mixing of different cell subtypes, displaying very little distinct clustering and no observable differentiation patterns. Diffusion Maps, however, shows both clustering of cell subtypes and trajectories that may correspond to differentiating cells (see below). T-SNE also shows the clearly defined clusters, with large gaps between some distinct groups. From these preliminary observations, we consider the t-SNE and Diffusion Maps embeddings for further examination.

**Figure 7:**
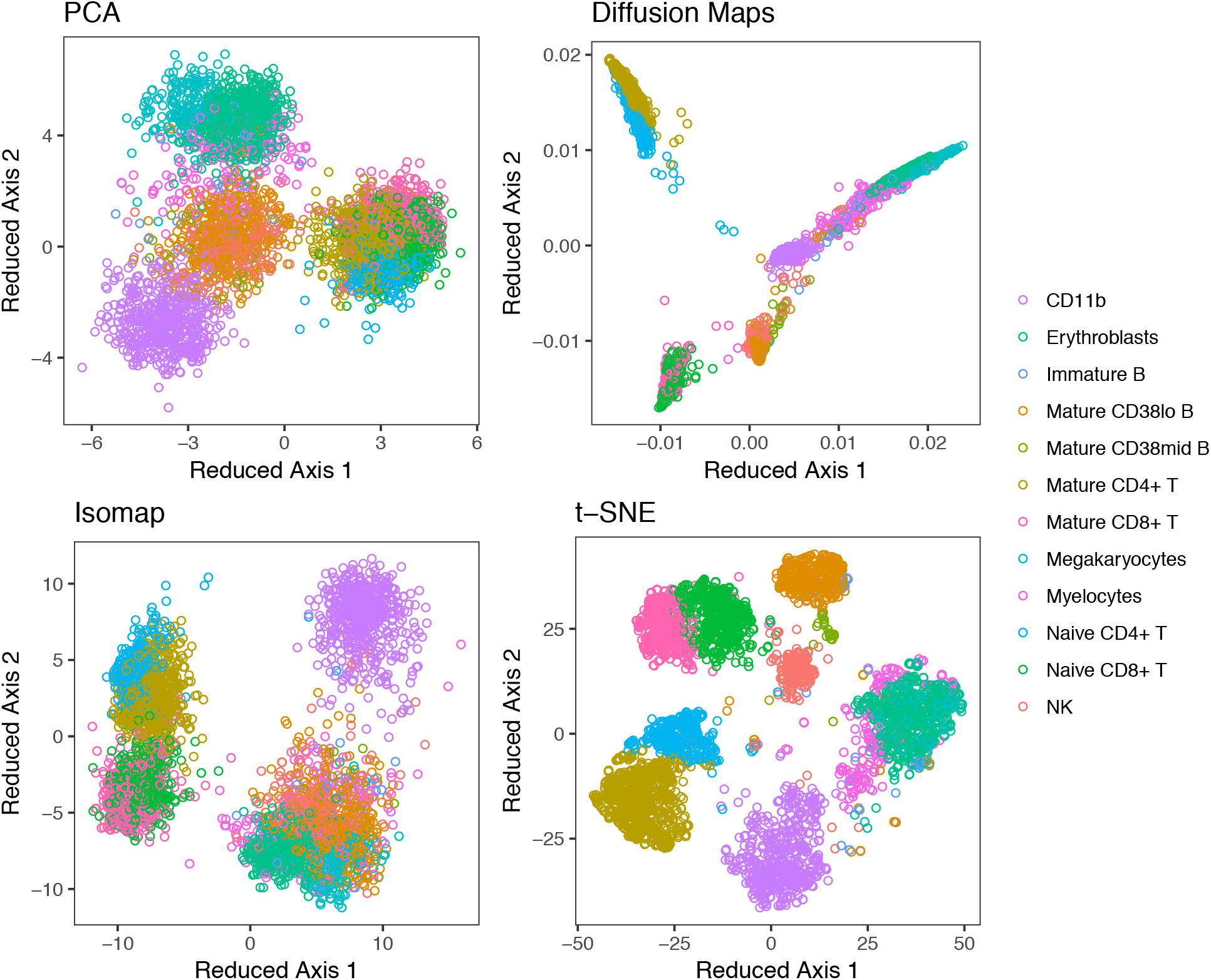
Two-dimensional embeddings of a random sample of 10,000 cells from the benchmark dataset for the four dimension reduction techniques. Manually gated cell subtypes are labeled.

Indeed, if we view the embedding results for just cells of the common lymphoid progenitor (CLP) lineage, which include natural killer (NK) cells, naive and mature CD8+, naive and mature CD4+, and immature, mature CD38mid, and mature CD38lo B cells, for all the four methods (Figure 8(a)), we observe that while all reductions show naive and mature CD8+ (and, CD4+) cells near each other (respectively), the greatest separation of the naive and mature subsets, as well as the CD4+ and CD8+ cells is, achieved by t-SNE and Diffusion Maps (Figure 8(a) and (b) top panels of (i) and (ii)). CD38 is a marker of developing B cells, with a loss of expression as B cells mature [6]. The differentiation trajectory from immature B, to CD38 mid, to CD38 lo B cells is observable in the Diffusion Maps projection (Figure 8(b)(i) bottom panel), but much less so for t-SNE (Figure 8(b)(ii) bottom panel). T-SNE thus shows an enhanced ability over the other methods to distinguish different cell subtypes, whereas Diffusion Maps shows an enhanced ability to distinguish differentiation trajectories.

**Figure 8:**
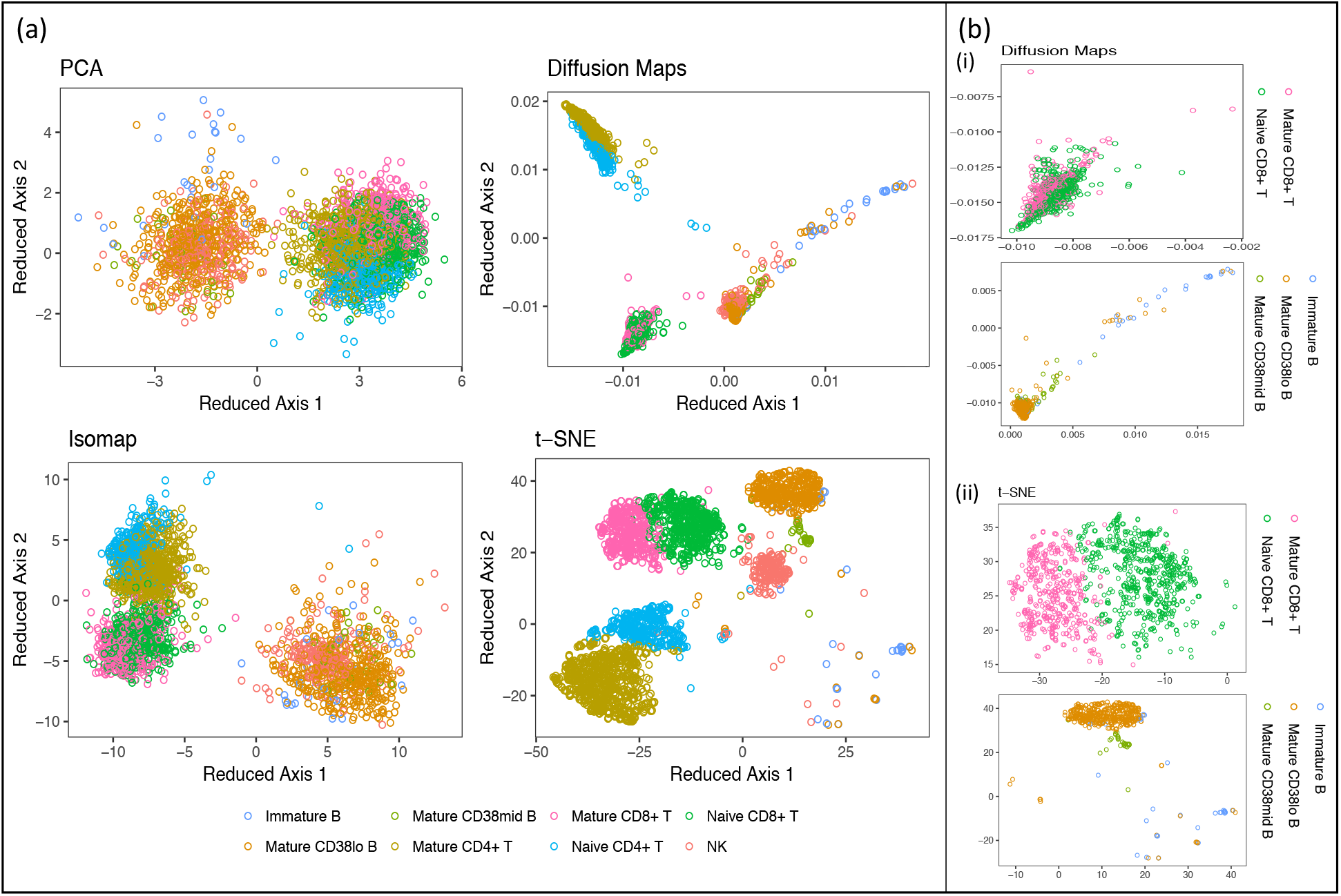
Common lymphoid progenitor (CLP) populations present in the two-dimensional embeddings of the four dimension reduction methods. (a) CLP populations for each method and (b) subsets of naive and mature CD8+ T cells and immature, mature CD38mid and mature CD38lo B cells for (i) Diffusion Maps and (ii) t-SNE.

## 4 Discussion

We have compared several popular dimension reduction techniques for their utility in information of interest from data generated by CyTOF. We chose to focus on these techniques since they make minimal assumptions on the nature of relationships between the input cells, and thus can be used on datasets comprising both differentiating cells and cells of different lineages. We used several measures for the quality and utility of a reduction. Computation time is a relevant measure since CyTOF experiments are often performed on up to hundreds of thousands of cells, and thus computation time may become a challenge using certain methods, as the end-users of these analyses are often experimental biologists and immunologists, who may not have access to high-throughput computing facilities. We developed a supervised heuristic algorithm, Neighborhood Proportion Error (NPE), to help answer the question of how well each technique preserves the empirical distribution of like neighbor frequencies amongst cell subtypes. We consider this an important property of the data that serves as a proxy of the relative clustering of manually gated cell sub-types, and one that an employed dimension reduction algorithm should ideally seek to preserve. We sought out similar information via visualization of the two-dimensional reductions vis-a-vis the manually gated cell types, which also provided visual information as to whether the methods preserved differentiation trajectories. Finally, we used residual variance to measure the global total information loss with each dimension reduction method. Isomap performed poorly on all measures except for RV, and PCA performed poorly on all measures except computation time and RV. Since preservation of relationships between and within known cell subtypes is of import in a dimension reduction, which was measured by the NPE and visualization, we consider Isomap and PCA poor performers for CyTOF, despite the good RV results. Diffusion Maps and t-SNE were the second-best performers for computation time, after PCA, and Diffusion Maps was able to clearly separate known cell subtypes, and identified differentiation trajectories. T-SNE was able to separate known cell types more thoroughly than Diffusion Maps, but was not necessarily able to capture differentiation trajectories, and does not provide information regarding the strength of similarity between different clusters, as was also noted in [5, 46]. While Becher et al. [5] and Wong et al. [46] used Isomap to determine relationships between cell subtypes, we recommend using Diffusion Maps for this purpose. Indeed, Haghverdi et al. [20] found that for qPCR data of mouse hematopoietic and progenitor stem cells, as well as qPCR data of mouse cells in early stages of embryonic development, Diffusion Maps showed clearer differentiation trajectories than Isomap. We thus recommend the complementary approach of using t-SNE and Diffusion Maps on CyTOF data. If this data contains cell types of heterogeneous lineages, each method will be able to contribute different information that in concert will allow researchers to obtain a better understanding of both the different cell types and potential differentiation lineages comprising the population.

## Supporting information

## Acknowledgments

This research was supported by the National Science Foundation - Division of Mathematical Sciences Award 1460967. Additionally, A.K. acknowledges support from the National Cancer Institute of the National Institutes of Health postdoctoral fellowship award F32CA214030.

