## Supplementary material for "Comparative Analysis of Linear and Nonlinear Dimension Reduction Techniques on Mass Cytometry Data"

#### S1 Benchmark Datasets

The benchmark dataset used in the main text for the comparative analysis is originally from Bendall et al. (1), and consists of a total of 167,044 bone marrow mononuclear cells (BMMCs) derived from healthy primary bone marrow. CyTOF was used to measure the intensity of 13 markers, and the data was manually gated into 24 populations (Table S1). An additional benchmark dataset of BMMC cells from Levine et al. (2) was analyzed using the same comparison metrics. CyTOF was used to measure the intensity of 32 markers for this dataset, and it was manually gated into 14 cell populations (Table S2). Note that for the 32 marker dataset, the counts stated in (3) include duplicated entries in the unassigned counts for the labeled cells. We have removed the duplicated entries in our analysis.

#### S2 Parameters for the dimension reduction techniques

We aimed to use parameters suggested by the authors of the respective methods and/or respective implementations in R (Table S3). For choosing the Isomap neighborhood size, $k$, we chose a range of $k$ $(5, 10, 15, 20)$ and performed the Isomap reduction to dimension $n = 2$ of the 13 marker benchmark dataset, subset 1 used in the main text (Figure S1) to identify the $k$ with the lowest residual variance.

Table S1: The frequency and % of labeled cells from the 13 marker dataset for each manually gated cell population is shown. For visualization purposes (Section 3.4), certain subpopulations were grouped together, while other very low frequency populations were not visualized, and are indicated as removed ($R$), under the Classification heading.

| **Population** | **Frequency** | **% Labeled Cells** | **Classification** |
|---|---|---|---|
| CD11b- Monocyte cells | 912 | 1.12 | CD11b |
| CD11bhi Monocyte cells | 6779 | 8.29 | CD11b |
| CD11bmid Monocyte cells | 1278 | 1.56 | CD11b |
| CMP cells | 253 | 0.31 | $R$ |
| Erythroblast cells | 12030 | 14.72 | |
| GMP cells | 73 | 0.09 | $R$ |
| HSC cells | 261 | 0.32 | $R$ |
| Immature B cells | 502 | 0.61 | |
| Mature CD38lo B cells | 7796 | 9.54 | Mature B Cells |
| Mature CD38mid B cells | 608 | 0.74 | Mature B Cells |
| Mature CD4+ T cells | 13964 | 17.08 | |
| Mature CD8+ T cells | 7821 | 9.57 | |
| Megakaryocyte cells | 3684 | 4.51 | |
| MEP cells | 194 | 0.24 | $R$ |
| MPP cells | 152 | 0.19 | $R$ |
| Myelocyte cells | 3025 | 3.70 | |
| Naive CD4+ T cells | 6987 | 8.55 | |
| Naive CD8+ T cells | 9564 | 11.70 | |
| NK cells | 3864 | 4.73 | |
| Plasma cell cells | 468 | 0.57 | $R$ |
| Plasmacytoid DC cells | 293 | 0.36 | $R$ |
| Platelet cells | 5 | 0.01 | $R$ |
| Pre-B I cells | 240 | 0.29 | $R$ |
| Pre-B II cells | 994 | 1.22 | $R$ |
| Total Labeled | 81747 | 100 | |
| Unassigned | 85297 | 0 | R |

Table S2: The frequency and % labeled cells from the 32 marker dataset for each manually gated cell population is shown. For visualization purposes (Section S4), certain subpopulations were grouped together, while other very low frequency populations were not visualized, and are indicated as removed ($R$), under the Classification heading.

| **Population Name** | **Frequency** | **% Labeled Cells** | **Classification** |
|---|---|---|---|
| Basophils | 1207 | 1.16 | |
| CD16- NK cells | 3905 | 3.75 | NK cells |
| CD16+ NK cells | 2248 | 2.16 | NK cells |
| CD34+CD38+CD123- HSPCs | 3295 | 3.16 | HSCs |
| CD34+CD38+CD123+ HSPCs | 304 | 0.29 | HSCs |
| CD34+CD38lo HSCs | 916 | 0.88 | HSCs |
| CD4 T cells | 26366 | 25.31 | |
| CD8 T cells | 20108 | 19.30 | |
| Mature B cells | 16520 | 15.86 | |
| Monocytes | 21099 | 20.25 | |
| pDCs | 1238 | 1.19 | |
| Plasma B cells | 330 | 0.32 | $R$ |
| Pre B cells | 6135 | 5.89 | |
| Pro B cells | 513 | 0.49 | $R$ |
| Total Labeled | 104184 | 100 | |
| Unassigned | 57259 | 0 | $R$ |

Table S3: Parameters

| Method | Parameters | Justification |
|---|---|---|
| Isomap | Neighborhood size ($k$) = 5 | A residual variance plot for different $k \in \{5, 10, 15, 20\}$ of the Isomap algorithm applied to the 13 marker dataset used in the main text with output dimension $n = 2$ shows that the lowest residual variance is found when $k = 5$ (Figure S1) (4). |
| t-SNE | $\theta = 0.5$ (default) | Rtsne package default (5), based on optimized threshold estimate for a trade-off between speed and accuracy in a Barnes-Hut implementation of t-SNE (6). |
| | Perplexity = 30 (default) | Rtsne package default (5), based on estimate in (7). |
| Diffusion Maps | Gaussian kernel width ($\sigma$) = default local estimate ('local') | A $\sigma$ for each data point is calculated in proportion to the estimated dimensionality at the local position (8; 9). |

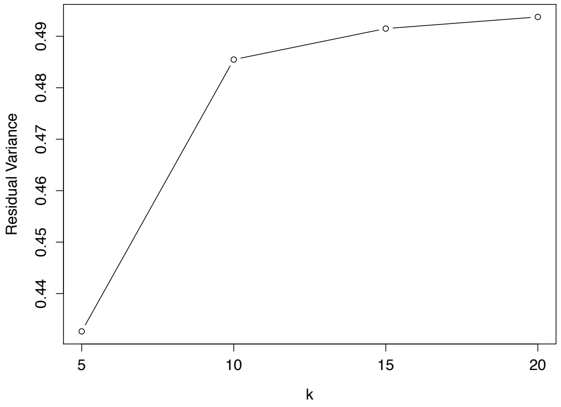


Figure S1: Residual variance of the Isomap dimension-reduced ($n = 2$) 13 marker dataset (subset 1) for different neighborhood sizes ($k$).

### S3 Visualization results for the 13 marker benchmark datasets: subsets 2 and 3

Taking different 10,000 cell subsets of the 13 marker benchmark dataset produces no qualitative differences in the visualizations (Figures S2 and S3).

### S4 Analysis results for the 32 marker benchmark dataset

Using the dataset from Levine et al. (2) (described in Section S1) as the benchmark dataset produces results qualitatively similar to the Bendall et al. (1) dataset analyzed in the main text (Figures S4 and S5).

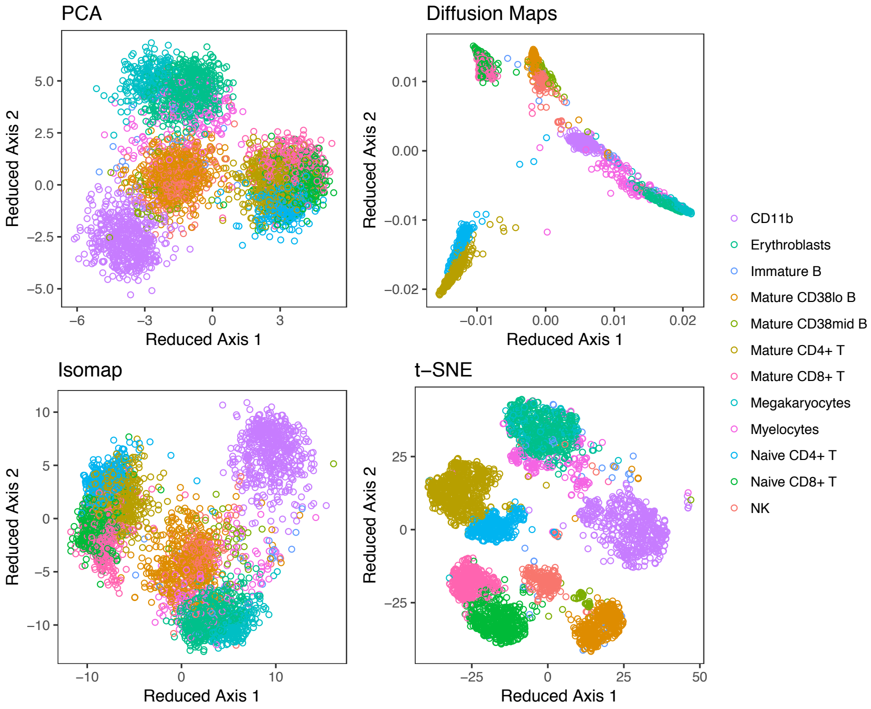


Figure S2: Two-dimensional embeddings of a different random sample from the main text (subsample ‘10k2’) of 10,000 cells from the benchmark dataset for the four dimension reduction techniques. Manually gated cell subtypes are labeled.

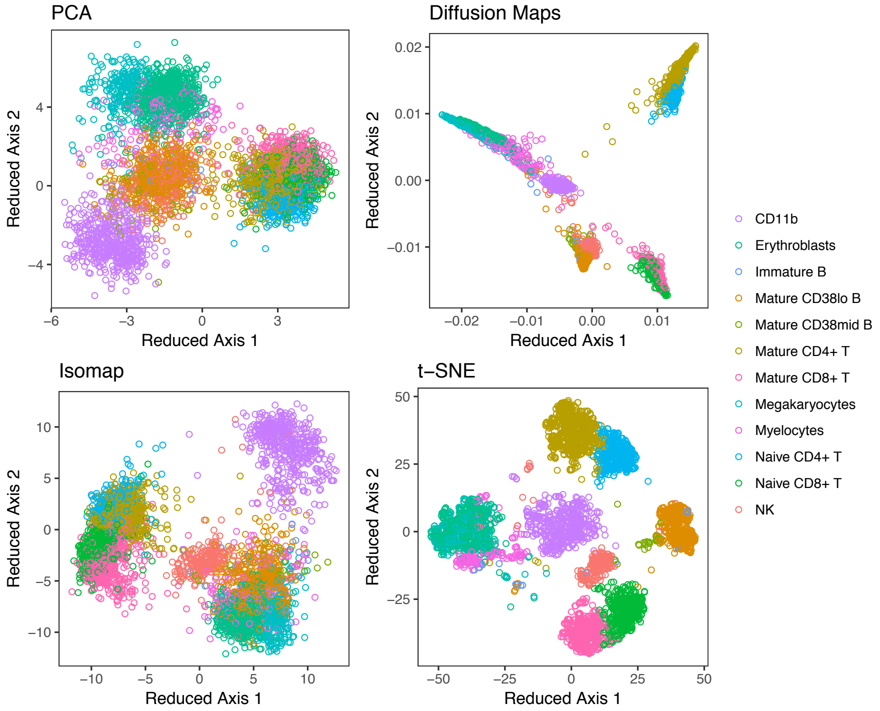


Figure S3: Two-dimensional embeddings of a different random sample from the main text (subsample ‘10k3’) of 10,000 cells from the benchmark dataset for the four dimension reduction techniques. Manually gated cell subtypes are labeled.

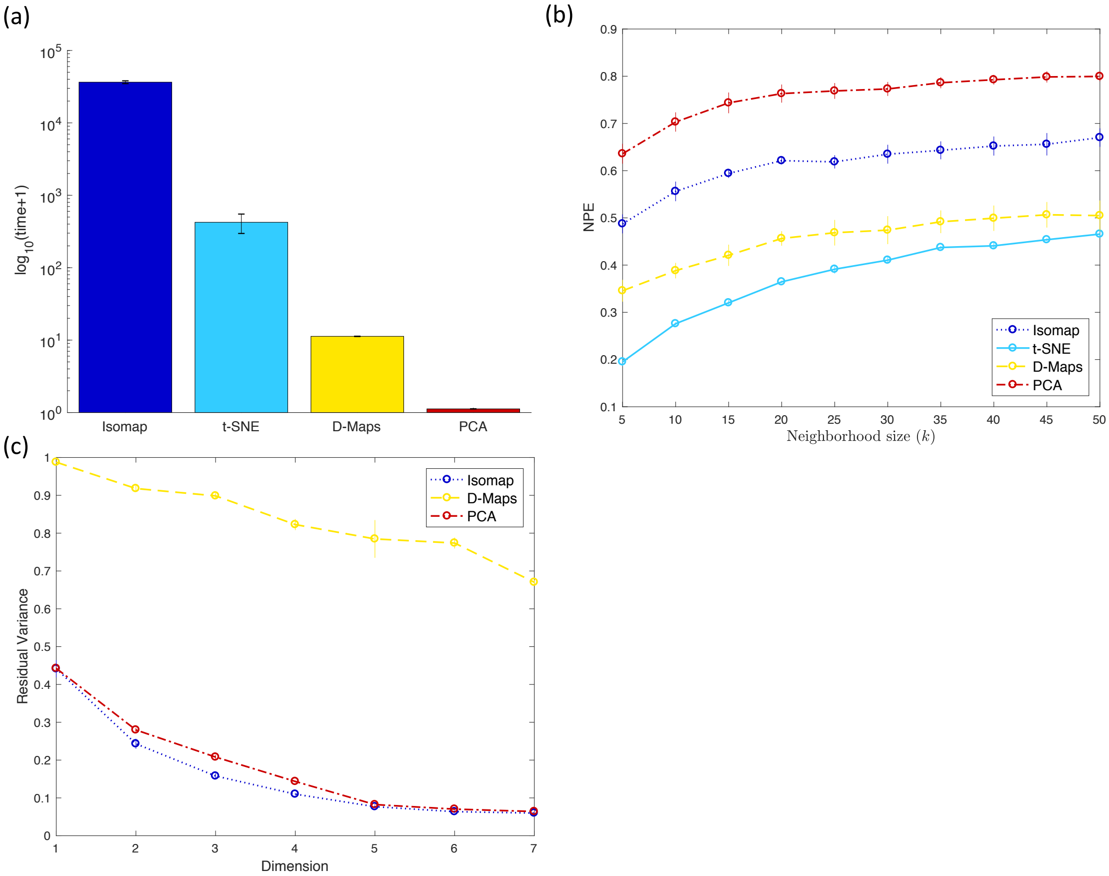


Figure S4: Comparative analysis of the 32 marker benchmark dataset with respect to (a) computation time, (b) neighborhood proportion error (NPE), and (c) residual variance.

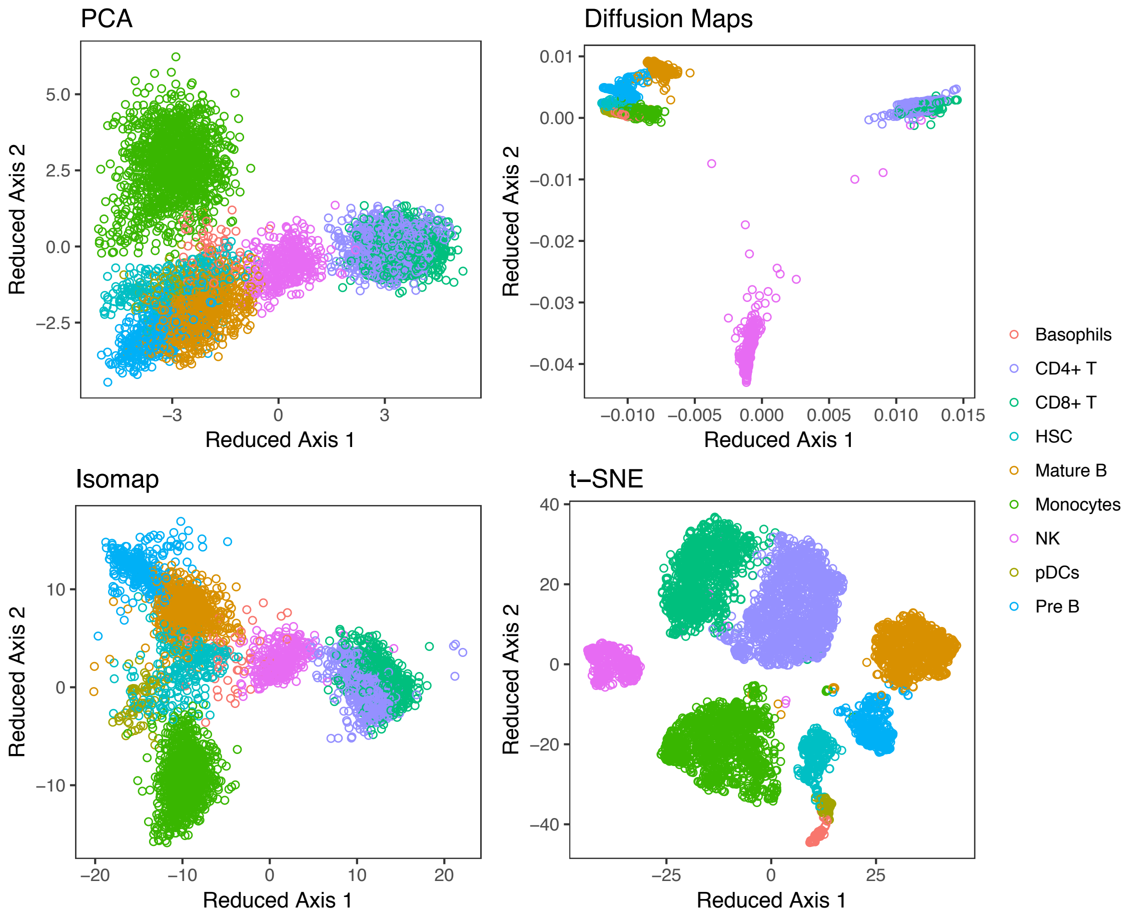


Figure S5: Two-dimensional embeddings of a random sample of 10,000 cells from the 32 marker benchmark dataset for the four dimension reduction techniques. Manually gated cell subtypes are labeled.
